# Unraveling Integration-Segregation Imbalances in Schizophrenia Through Topological High-Order Functional Connectivity

**DOI:** 10.1101/2024.10.03.616506

**Authors:** Qiang Li, Wei Huang, Chen Qiao, Huafu Chen

## Abstract

**Background:** The occurrence of brain disorders correlates with detectable dysfunctions in the specialization of brain connectomics. While extensive research has explored this relationship, there is a lack of studies specifically examining the statistical correlation between the integration and segregation of psychotic brain networks using high-order networks, given the limitations of low-order networks. Moreover, these dysfunctions are believed to be linked to information imbalances in brain functions. However, our understanding of how these imbalances give rise to specific psychotic symptoms remains limited.

**Methods:** This study aims to address this gap by investigating variations at the topological high-order level of the system with regard to specialization in both healthy individuals and those diagnosed with schizophrenia. By employing graph-theoretic brain network analysis, we systematically examine information integration and segregation to delineate system-level differences in the connectivity patterns of brain networks.

**Results:** The findings indicate that topological high-order functional connectomics highlight differences in the connectome between healthy controls and schizophrenia, demonstrating increased cingulo-opercular task control and salience interactions, while the interaction between subcortical and default mode networks, dorsal attention and sensory/somatomotor mouth decreases in schizophrenia. Furthermore, we observed a reduction in the segregation of brain systems in healthy controls compared to individuals with schizophrenia, which means the balance between segregation and integration of brain networks is disrupted in schizophrenia, suggesting that restoring this balance may aid in the treatment of the disorder. Additionally, the increased segregation and decreased integration of brain systems in schizophrenia patients compared to healthy controls may serve as a novel indicator for early schizophrenia diagnosis.

**Conclusion:** We discovered that topological high-order functional connectivity highlights brain network interactions compared to low-order functional connectivity. Furthermore, we observed alterations in specific brain regions associated with schizophrenia, as well as changes in brain network information integration and segregation in individuals with schizophrenia.

## 1. INTRODUCTION

Understanding the intricate neural dynamics associated with schizophrenia is a complex endeavor that has garnered significant attention to help us better cure schizophrenia. One crucial aspect of unraveling the underlying mechanisms of this debilitating psychiatric disorder lies in the examination of functional connectivity with functional magnetic resonance imaging (fMRI) [1]–[3]. In regards to schizophrenia, irregularities in functional connectivity have gained significant attention in our endeavor to comprehensively understand the neurobiological underpinnings of the disorder [4]–[7].

When examining the human brain as an intricately non-linear dynamic system, explaining the mechanisms of information integration and segregation becomes pivotal [8]–[12]. Understanding how information is transmitted between brain states is crucial, as poor information integration and segregation may cause mental illnesses [5], [13]. As a result, tracking and quantifying informational interactions between controls and people with schizophrenia has the potential to dramatically improve our understanding of the fundamental mechanisms underlying this illness. A normal brain functions effectively by segregating and integrating regular information [14], [15]. Maintaining a delicate balance between these activities is critical, as any disruption in this equilibrium could result in severe brain illnesses, including schizophrenia [16]. Detecting deviations from this balance becomes imperative for understanding schizophrenia, and one potential avenue for achieving this understanding involves quantifying the levels of information segregation and integration within the brain.

Exploring the brain from a functional connectivity perspective always tends to measure the topology of brain network properties, and many studies have investigated the similarities and differences between healthy controls and schizophrenia [3], [17]. Certain brain networks have been found to be altered in schizophrenia, but little research has examined the relationship between the disorder and high-order networks, which provide a more comprehensive view of complex systems by capturing interactions among multiple nodes simultaneously rather than just pairwise connections. This is because low-order networks may overlook beyond pairwise information, and since the brain is a complex system, information interactions in the brain typically extend beyond pairwise connections. Meanwhile, information interaction and segregation play a super-significant role in maintaining normal brain function, and indeed, many brain diseases will more or less disrupt this information interaction balance [5], [13]. Therefore, this study delves into the altered patterns of functional connectivity segregation in schizophrenia using high-order networks, shedding light on how altered brain network information interaction may contribute to the manifestation and detection of some biomarkers in schizophrenia.

## 2. THEORETICAL ANALYSIS

### 2.1. Participants

The Centers of Biomedical Research Excellence (COBRE) respiratory data includes both raw structural and fMRI data, comprising schizophrenia (SZ) and healthy controls (HC) [1], [18]. To ensure data integrity, all subjects underwent thorough screening, and those with a history of neurological disorders, severe head trauma resulting in more than 5 minutes of loss of consciousness, or a history of substance dependence or abuse within the last 12 months were excluded from the study.

### 2.2. Image Acquisition

In the group of COBRE subjects, a multi-echo MPRAGE (MEMPR) sequence was used with the following parameters: TR/TE/TI = 2530/[1.64, 3.5, 5.36, 7.22, 9.08] ms / 900 ms, flip angle = 7°, FOV = 256 mm × 256 mm, slab thickness = 176 mm, matrix = 256 × 256 × 176, voxel size = 1 mm × 1 mm × 1 mm, number of echoes = 5, pixel bandwidth = 650 Hz, total scan time = 6 minutes. With five echoes, the TR, TI, and partition encoding times for the MEMPR are similar to those of a conventional MPRAGE, resulting in comparable GM/WM/CSF contrast.

The resting-state fMRI data were acquired through echo-planar imaging (EPI) with ramp sampling correction, utilizing the intercommissural line (AC-PC) as a reference (TR = 2 s, TE = 29 ms, matrix size = 64 × 64, 32 slices, voxel size = 3×3×4 *mm*^3^). More details about scanning parameters can be found in [1], [18]. In this study, we opted to include 54 individuals diagnosed with SZ and 70 HC. The age range for individuals in each group spans from 18 to 65 years, and no significant differences in age were observed between the two groups (p=0.27).

### 2.3. Image Preprocessing

The standard pre-processing steps were conducted using the Statistical Parametric Mapping (SPM8) package from the Wellcome Institute of Cognitive Neurology, London, such as slice timing, realign, coregister, normalize, denoise, smooth, and multiple regressions were utilized on the time series to eliminate the following nuisance effects: linear trend, 6 motion parameters along with their temporal derivatives, the quadratic terms of these 12 parameters, 5 components derived from PCA of CSF, and 5 components derived from PCA of WM, and then fMRI signal was filtered with a bandpass of [0.01, 0.1]Hz. More detailed information on the preprocessing steps can be found in [20].

### 2.4. Functional Connectivity

#### Static Low-order Functional Connectivity

The time courses were extracted from 10mm-diameter spheres based on the 263 sets of coordinates identified by [19], and for an fMRI signal of 𝒩 networks from power-based brain parcellation, *x*(*i*) (1 ≤ *i* ≤ 𝒩, 𝒩 =263), informally, given 𝒩 =(*x*_1_, *x*_2_, *x*_3_,…, *x*_*𝒩*_) points with a constant time interval *t*. A cross-correlation matrix of Pearson values was obtained from the time series data, and Fisher’s z transformation was applied to each of the 263 brain regions along with every other region, as depicted in Fig.1. The low-order functional connectivity, as measured using traditional Pearson correlation, which is present as

**Fig. 1:**
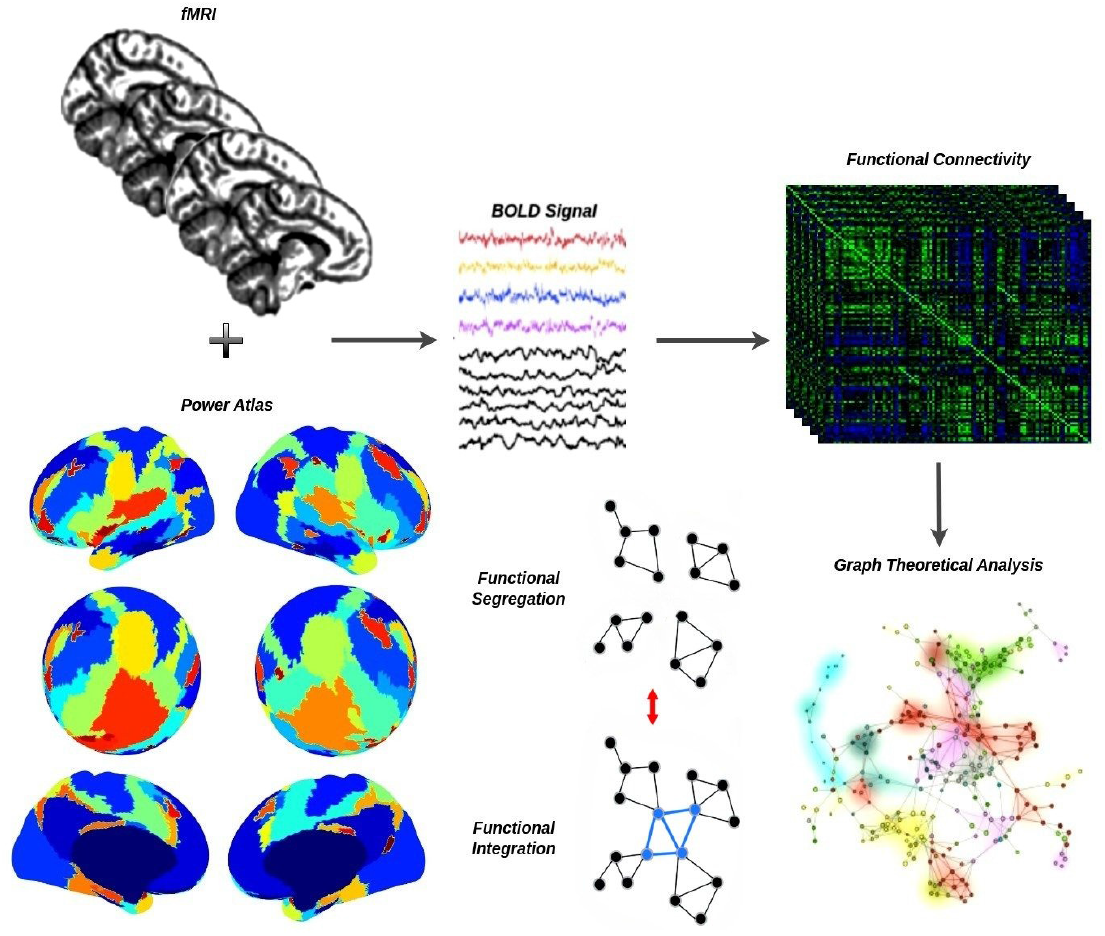
A flowchart for the construction of a functional brain network by preprocessed fMRI. First, time series extraction from preprocessed fMRI data within each functional unit (i.e., network node). Second, estimate functional connectivity. Third, visualization of functional connectivity as tree graphs (i.e., network edges and network nodes). Here, the Power Atlas was presented on an inflated and sphered brain surface, and it contains 264 nodes and can basically be categorized into 14 brain sub-networks, which indicates a difference in color [19].

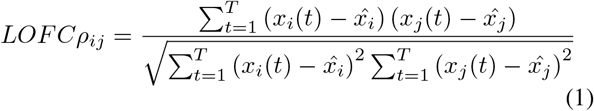

where 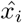 and 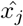 are the average of the rsfMRI signals at regions *i* and *j*.

For each participant, assigned edge weights were determined by ranking these scores individually, and subsequently, these values were standardized and centered across the entire participant group. Here, we totally identify 14 intrinsic functional subnetworks of the human brain: Sensory/Somatomotor Hand (SMH), Sensory/Somatomotor Mouth (SMM), Cingulo-Opercular Task Control (CO), Auditory (AU), Default Mode (DM), Memory Retrieval (MR), Visual (VS), Fronto-Parietal Task Control (FP), Salience (SA), Subcortical (SC), Ventral Attention (VA), Dorsal Attention (DA), Cerebellar (CB), and unknown (UN) networks.

### 2.5. High-order Functional Connectivity from the Correlation of Correlation

According to the proposal by Han et al. [21], the pairwise connection represented by topological high-order functional connectivity (tHOFC) differs from that of low-order functional connectivity due to variations in input signals. Nonetheless, tHOFC characterizes the interaction between two brain regions, establishing high-order functional connectivity. The LOFC are derived from the rsfMRI time series, but these time series serve as local LOFC profiles for tHOFC. Notably, a significant variation exists between LOFC and tHOFC for the same two brain regions, attributed to differences in input signals [21], [22].

#### 2.5.1 Topological High-order Functional Connectivity

Letting *ρ*_*i*_ represents the LOFC profile for region *i*, and {*ρ*_*i*·_ = *ρ*_*ik*|*k*∈ℝ,*k*≠ *i*_}. Then the tHOFC can be denoted as,

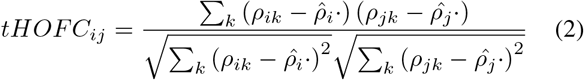

The tHOFC can be thought of as a “correlation of correlation,” and it can highlight interaction information in the brain and then supply more rich information to detect the difference between HC and SZ, as described in Fig.3.

### 2.6. Comparative Classification Performance Using tHOFC and LOFC

#### Divisive Normalization

Divisive normalization (DN) is a phenomenological model that captures the nonlinear response properties observed widely across sensory cortical areas [23]. It involves scaling the response of a neuron by the average activity of a group of neurons or by the overall level of activity in the neural population. Essentially, the activity of a neuron is divided by a measure of the collective activity of its neighbors or the population, thereby *normalizing* its response [23]. Optimizing with the incorporation of more hyperparameters becomes increasingly challenging. In our case, we adapted and simplified DN [24] and applied it to enhance features in functional connectivity matrices. This can be mathematically represented as follows:

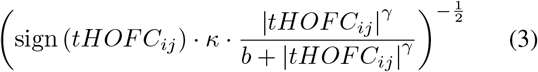

There are several hyperparameters (i.e., *κ, γ, b*), and after tuning parameters, we selected *γ*=2, *b* = 0.0625, and *κ*=0.5. Here *tHOFC*_*ij*_ refers to tHOFC matrix.

#### Machine Learning

The Support Vector Machine (SVM) and K-Nearest Neighbors (KNN) algorithms [25] are employed to classify HC and SZ using functional connectivity as features. To assess model classification performance, we employed 10 rounds of 5-fold cross-validation, ensuring robust evaluation across multiple iterations. The SVM classifier is trained using a linear kernel function and standardized data to predict class labels on the test set. Similarly, a KNN classifier with 15 neighbors is trained and evaluated for comparison. Both classifiers’ evaluation metrics are computed and presented, providing insights into their effectiveness in distinguishing between healthy controls and patients based on functional connectivity patterns (tHOFC vs. LOFC).

#### Deep Nets

A popular pretrained convolutional neural network (CNN), known as AlexNet [26], which was developed in the field of state-of-the-art artificial intelligence, was used in this study for transfer learning and classification of HC and SZ. The architecture and training parameters for the transfer learning of pretrained CNNs are specified as follows: First, the layer graph was extracted from the pretrained network, and the list of layers was converted into a layer graph. Since the pretrained deep networks work with color images of fixed sizes and the input functional connectivity matrices were grayscale images, we converted them into RGB images by stacking three copies of the grayscale image to create a corresponding 3-D RGB array. The images were then resized to match the input size required by the deep network. In pretrained networks, the final layer with learnable weights typically consists of a fully connected layer. This fully connected layer was substituted with a new fully connected layer designed for the number of classes in the new dataset, which in this study is 2, as illustrated in Fig.2. For training, stochastic gradient descent with a momentum optimizer was employed, with a momentum value set to 0.9. The gradient threshold method used was based on the L2 norm, and the minimum batch size was set to 10. Training proceeded with a maximum of 54 epochs, using an initial learning rate of 0.0003 which remained constant throughout training. Prior to each training epoch, the training data were shuffled, and similarly, the validation data were shuffled before each validation session. Additionally, a factor of 0.0001 was applied for L2 regularization (weight decay).

**Fig. 2:**
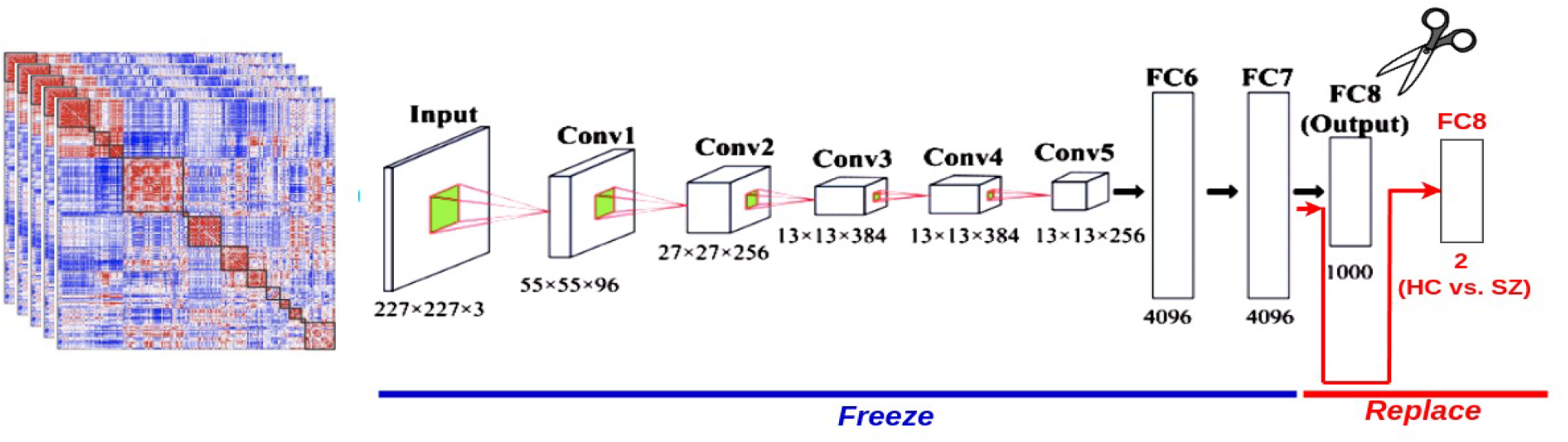
Deep nets for HC and SZ classification using LOFC and tHOFC. The deep nets for brain disorder classification utilize the frozen weights of AlexNet in previous layers, with only the last layer being replaced to classify into 2 classes instead of 1000.

#### Evaluation metrics

Let PL represent the total number of SZ functional connectivity matrices, NL represent the total number of HC functional connectivity matrices, TP represent the true positives (number of SZ functional connectivity matrices correctly identified as SZ), FP represent the false positives (number of HC functional connectivity matrices incorrectly identified as SZ), TN represent the true negatives (number of HC functional connectivity matrices correctly identified as HC), and FN represent the false negatives (number of SZ functional connectivity matrices incorrectly identified as HC).

- Accuracy

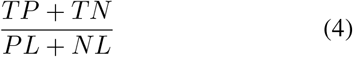

- Sensitivity

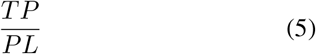

- Specificity

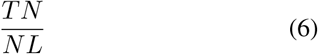

- Precision

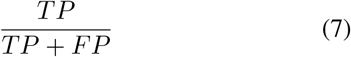

- F1-score

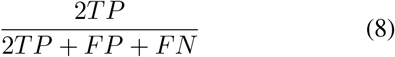

The receiver operating characteristic (ROC) curve is a probability curve created by plotting the true positive (TP) rate against the false positive (FP) rate at various threshold settings. The area under the ROC curve (AUC), which ranges from 0.5 to 1.0, represents the performance measure of a classifier. A higher AUC indicates that the classifier is better at distinguishing between HC and SZ subjects.

### 2.7. Brain System Segregation, Integration, and Their Balance in FC Networks

The analysis of brain networks through graph theory will provide brain network properties, elucidating the information interaction among the topological networks [8], [27], [28]. Here, we aim to quantify the extent of information segregation and integration in both healthy controls and individuals diagnosed with schizophrenia. Two measures of brain network segregation were evaluated here: (i) modularity and (ii) system integration and segregation.

#### Modularity

Modularity quantifies the degree to which the entire brain network can be divided into separate functional systems [29].

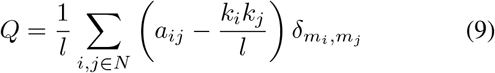

where *a*_*ij*_ is connection weights between *i* and *j. l* is the number of connections, *k*_*i*_ is the degree of node *i, m*_*i*_ is the module containing node *i*, and 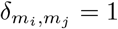 if 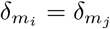 and otherwise. High modularity reflects a network in which regions are strongly grouped together based on their interactions, while these groups remain relatively isolated from one another. This can be seen as greater segregation, where the brain’s different functional systems are more distinct and specialized.

#### Brain Network Integration and Segregation

To quantify brain network integration and segregation, we used a second, more advanced approach to measure it. In order to estimate the brain information integration, we need to first decompose FC into eigenvectors *U* and eigenvalues Λ,

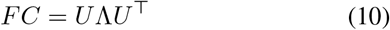

where a decreasing order of Λ was used to rearrange the eigenmodes. A few eigenvalues that were negative were set to zero. Then, we applied nested-spectral partition to detect hierarchical modules in the brain FC [30]. During this nested partitioning process, the modular number *M*_*i*_(*i* = 1,…, *N*) and the modular size *S*_*j*_ (*j* = 1, …, *M*_*i*_) were obtained. Then, the segregation between the modules and the integration within the modules may be measured at each level:

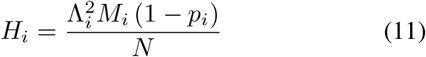

where *N* refers to normalizing the modular number, *M* refers to the range [0, 1], and *p*_*i*_ = Σ_*j*_|*S*_*j*_ − *N/M*_*i*_|*/N*, where N is a modular size correction factor that reflects the deviation from the optimized modular size in the *ith* level. Finally, the global integration can be calculated as follows:

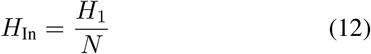

The total segregation component emerges from the several segregated levels:

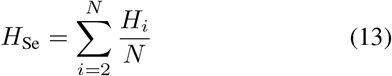

Here, a higher *H*_In_ or smaller *H*_Se_ reflects stronger network integration. Furthermore, to avoid artifacts for shorter fMRI series lengths in estimating information segregation, we applied the calibration process as suggested in [30], [31]. First, we measured individual information integration and segregation information, i.e., 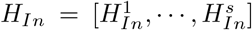, and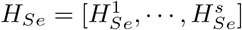., where *s* refers to the number of subjects. Second, the calibration results for each subject will be: 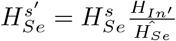 and 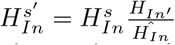. Here, 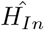 and 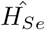 refer to group average integration and segregation individually, and 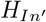 and 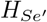 indicate integration and segregation from the average FC minus each individual integration and segregation score.

## 3 RESULTS

The static low-order and high-order functional connectivity of identified brain networks were estimated for both healthy controls and individuals with schizophrenia, with their differences illustrated in Fig.3. Initially, we observed that the tHOFC distinctly highlights more functional connectivity features compared to LOFC. The tHOFC not only provides us with a richer set of informative connectivity patterns but also maintains its fundamental topological structure. Specifically, tHOFC exhibits a robust network of interactions among cortical networks, and it may reveal nuanced relationships that contribute significantly to understanding the dynamic information interplay between various brain networks in brain. Consequently, we strongly recommend the application of tHOFC rather than LOFC in fMRI functional connectivity analysis. This approach promises to provide richer feature sets in functional connectivity matrices and potentially enhance the accuracy of schizophrenia diagnosis.

**Fig. 3:**
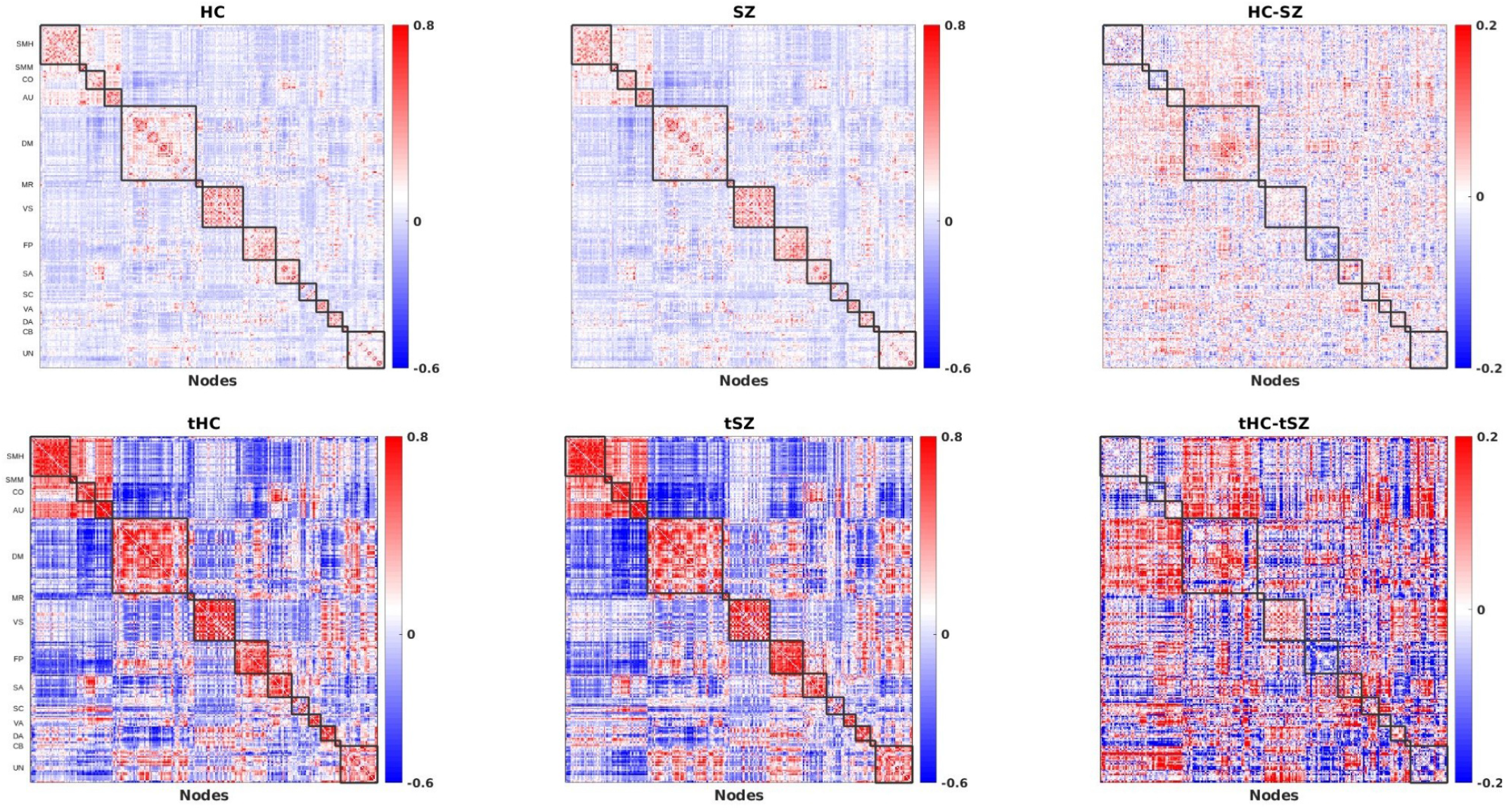
Psychosis functional brain connectomes from the COBRE dataset. The functional connectivity of healthy controls and individuals with schizophrenia was estimated using Pearson correlation based on the Power-Brain Networks Atlas [19]. Related brain regions were grouped into specific communities (here are 14 communities/brain networks, i.e., SMH, SMM, CO, AU, DM, MR, VS, FP, SA, SC, VA, DA, CB, UN). The top row shows the low-order functional connection and the difference between HC and SZ. The bottom row shows topological high-order functional connectivity based on topology connection information.

To explore the integration and segregation of topological information in HC and individuals with SZ, we transformed functional connectivities into a graph layout. This allowed us to visualize their interactions, as depicted in Fig.4A. Notably, compared to the functional connectivity matrices shown in Fig.3, HC and SZ exhibit distinct topological distribution shapes. Secondly, we observed that information tends to cluster within systems. Additionally, to further quantify this, the modularity of the whole brain network and each subnet was measured. Our findings suggest that individuals with SZ tend to show lesser segregation and more integration in subnets compared to HC subjects, specifically in the CO (p=0.043, t=2.436) and SC (p=0.028, t=2.221) subnets, as shown in Fig.4B. This indicates increased coordination between different brain regions in SZ, suggesting enhanced synchronized or coherent connectivity compared to HC in those subnets. Moreover, we show that tHOFC is more sensitive in capturing differences in brain network integration and segregation compared to LOFC. Additionally, to assess changes in functional connectivity strength, we plotted a functional connectogram, as presented in Fig.5. Consistently, this highlighted strong connectivity in the tHOFC compared to the LOFC, emphasizing significant differences that aid in uncovering abnormal connectivity patterns in the schizophrenia brain. In Fig.5, the strongest connections observed are between the cingulo-opercular task contro (CO) and salience (SA) networks, as well as between the salience (SA) and ventral attention (VA) networks. These brain networks are associated with attention, task control, and motivational behavior, and changes in their interactions can lead to instability in related behaviors [32]. Meanwhile, we also observed the strongest negative interactions between the default mode network (DM) and the subcortical (SC) network, as well as between the dorsal attention network (DA) and the sensory/somatomotor network (SMM). As a consequence, those increased or decreased interaction will change or destroy the information interaction balance in brain.

**Fig. 4:**
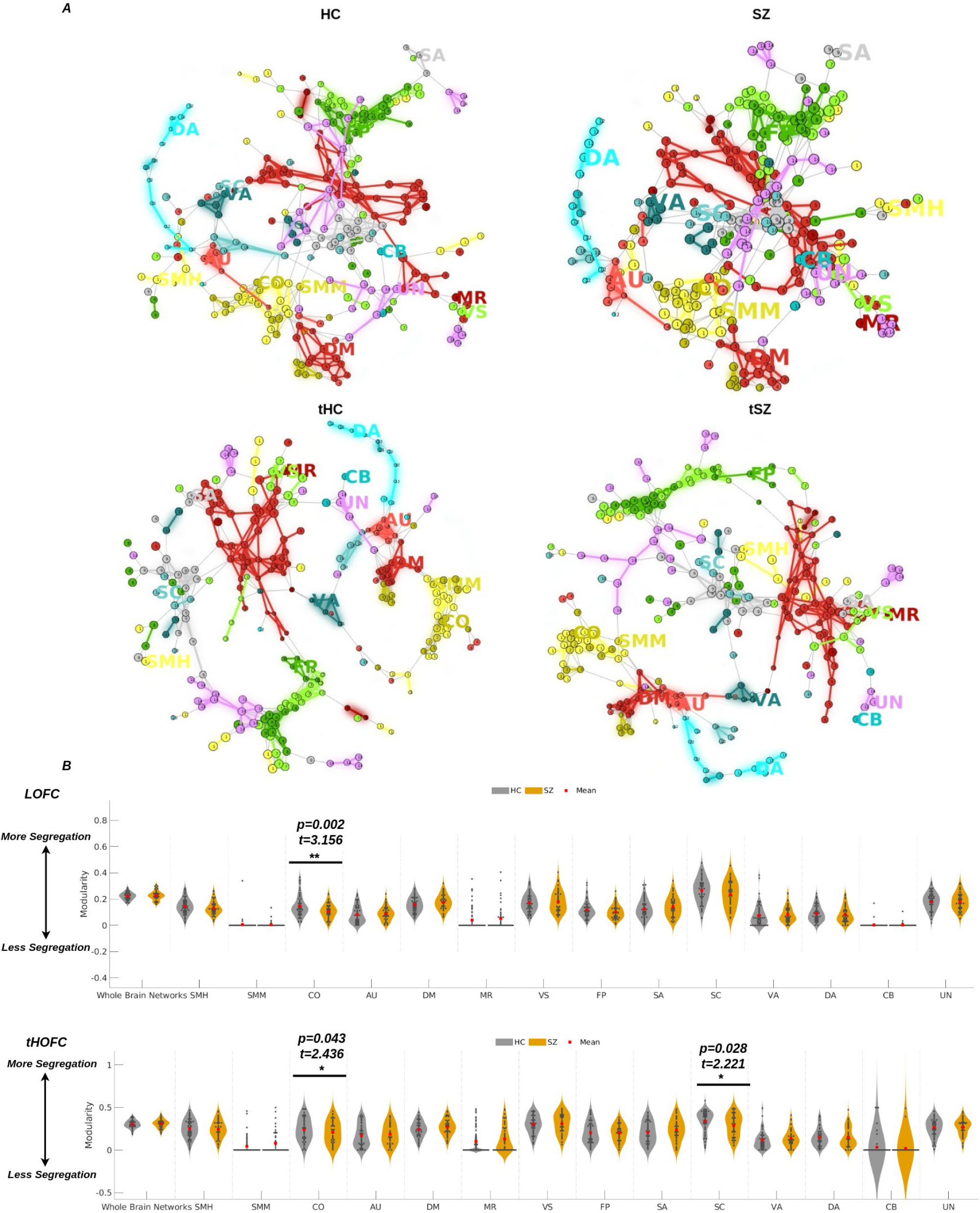
Topological functional brain networks in healthy and psychotic brains. The community-labeled low-order (LOFC) and high-order (tHOFC) functional connectograms for HC and SZ are shown individually in ***A***, where the 14 brain networks are labeled and used to investigate brain networks. The network interactions are visualized using a graph layout to assess their topological interactions in HC (first column, LOFC vs. tHOFC) and SZ (second column, LOFC vs. tHOFC). The associated quantities measured using modularity for HC and SZ in LOFC and tHOFC are shown in ***B*** (*, *p* < 0.05, **, *p* < 0.01).

**Fig. 5:**
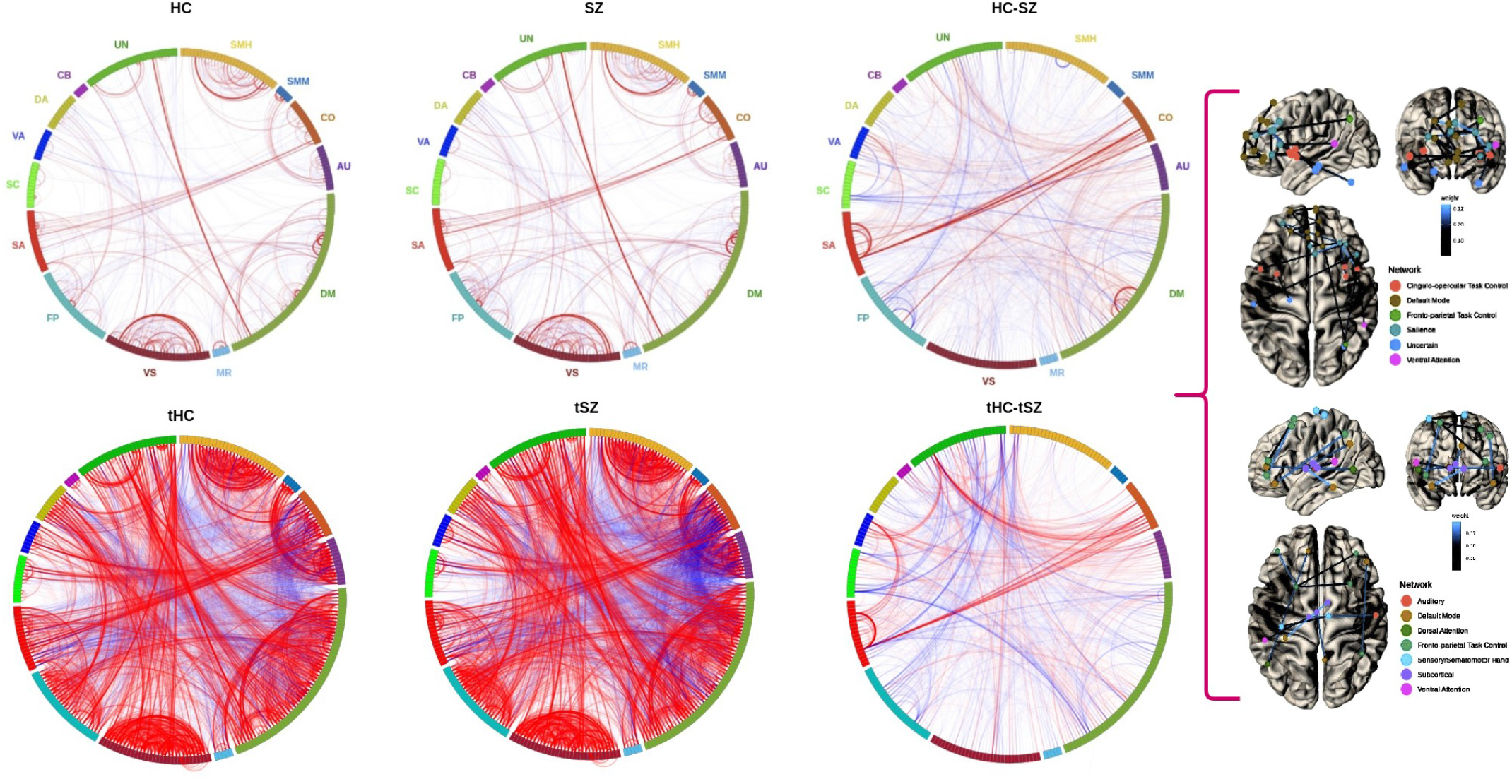
Functional connectograms were constructed for both HC and SZ groups. The left side illustrates the functional connectome for HC (LOFC vs. tHOFC), while the middle represents the functional connectome for SZ (LOFC vs. tHOFC). On the right side, the differential functional connectome between HC and SZ is depicted. Positive and negative interactions derived from these differences were projected onto the brain surface and visualized.

To further highlight the difference between tHOFC and LOFC, we fed both into machine learning and pretrained deep networks for classification. The evaluation metrics are presented in Tab.1. The results indicate that tHOFC performs significantly better than LOFC in classifying HC and SZ (i.e. for deep nets without DN, the accuracy improved from 67.66±2.4 to 92.37±1.5, AUC improved from 75.23±2.6 to 88.23±1.9; for deep nets with DN, the accuracy improved from 67.66±2.4 to 93.68±1.3, AUC improved from 75.23±2.6 to 91.35±0.5). This suggests that tHOFC provides richer features for deep networks compared to LOFC. Therefore, we strongly recommend using tHOFC instead of LOFC for future functional connectivity analyses.

**Table 1:** Classification performance comparison (average and standard errors, %) between the tHOFC and LOFC using different algorithms in the HC and SZ classifications. The top table shows classification scores using tHOFC, while the bottom table shows classification metrics using LOFC. The highest scores are labeled with bold colors.

| Categories | Accuracy | Sensitivity | Specificity | Precision | F1-score | AUC |
| --- | --- | --- | --- | --- | --- | --- |
| **tHOFc** |  |  |  |  |  |  |
| SVM | $84.23 \pm 1.2$ | $82.00 \pm 1.8$ | $85.29 \pm 2.2$ | $80.08 \pm 2.1$ | $80.00 \pm 1.3$ | $87.31 \pm 2.8$ |
| KNN | $71.83 \pm 1.2$ | $35.00 \pm 1.4$ | $100 \pm 0$ | $100 \pm 0$ | $46.45 \pm 1.1$ | $73.59 \pm 1.4$ |
| Deep Nets (AlexNet) | **$92.37 \pm 1.5$** | $84.43 \pm 2.2$ | $100 \pm 0$ | $100 \pm 0$ | $93.21 \pm 1.1$ | **$88.23 \pm 1.9$** |
| Deep Nets (AlexNet) + DN | **$93.68 \pm 1.3$** | $87.50 \pm 1.9$ | $100 \pm 0$ | $100 \pm 0$ | $95.70 \pm 1.3$ | **$91.35 \pm 0.5$** |
| **LOFC** |  |  |  |  |  |  |
| SVM | $78.89 \pm 2.6$ | $61.28 \pm 2.1$ | $93.07 \pm 1.3$ | $87.08 \pm 2.5$ | $72.73 \pm 2.3$ | $83.37 \pm 2.3$ |
| KNN | $74.89 \pm 0.3$ | $61.12 \pm 0.9$ | $86.08 \pm 0.4$ | $75.23 \pm 1.3$ | $67.57 \pm 2.3$ | $75.37 \pm 1.5$ |
| Deep Nets (AlexNet) | $67.66 \pm 2.4$ | $71.56 \pm 2.5$ | $61.23 \pm 2.3$ | $71.54 \pm 2.2$ | $72.67 \pm 2.5$ | $75.23 \pm 2.6$ |

Therefore, we utilized tHOFC to estimate integration and segregation scores, taking into account its prioritization in our analysis. To further quantify the information integration and segregation in schizophrenia, we applied Eq.12 and Eq.13 to quantify the integration and segregation scores in HC and SZ. We observed that there are a significant differences between HC and SZ, as shown in Fig.6, suggesting that the information balance is disrupted in schizophrenia. Meanwhile, our results are consistent with previous studies about information integration and segregation in HC and SZ, and they also found that information integration decreased and information segregation increased in SZ compared to HC [33].

**Fig. 6:**
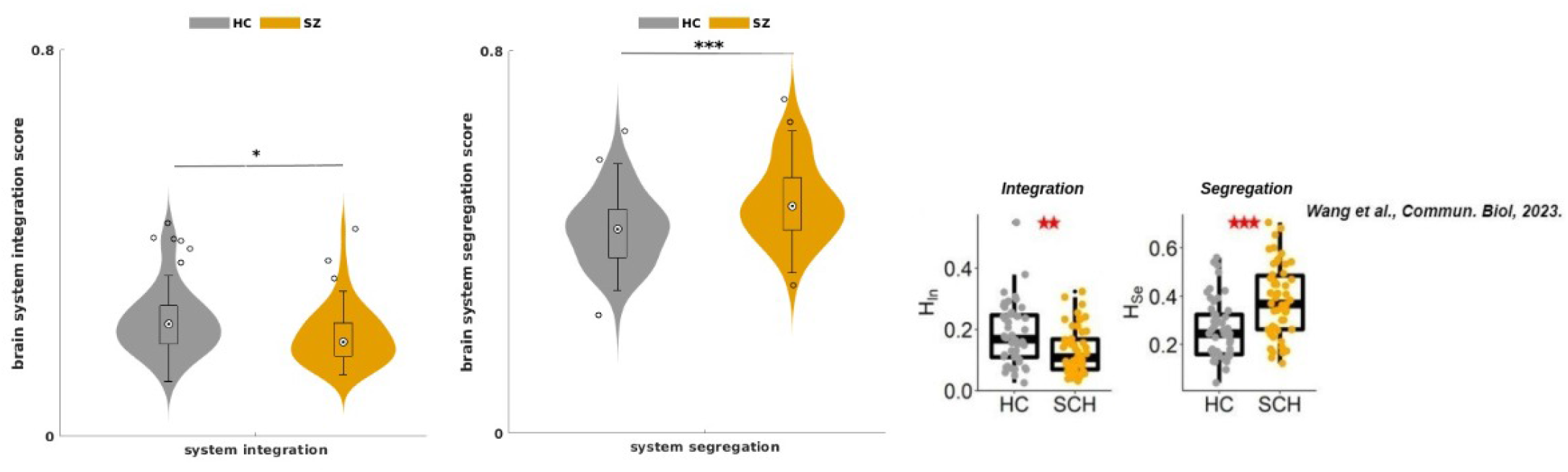
Brain system segregation and integration in psychosis brain. The brain system segregation score was measured; the information integration decreased (*, *p* < 0.05), but segregation increased in schizophrenia (***, *p* < 0.001). Meanwhile, we also found that previous studies supplied the same evidence that a significant brain system integration and segregation difference exists between healthy controls and schizophrenia (**, *p* < 0.01) [33].

In summary, we found that decreased segregation of brain systems in healthy controls relative to those who suffer from schizophrenia, and the other way around for information integration. Remarkably, individuals with schizophrenia exhibit an increase in the segregation of the brain system, which may provide a potential indicator for the treatment of schizophrenia. Furthermore, disrupting the brain system’s information balance may play a critical role in schizophrenia, and assessing information integration and segregation may provide a window into how we can early detect and treat schizophrenia.

## DISCUSSION

Examining the functional connectivity of psychosis is crucial for understanding mental illnesses. In our investigation, we observed abnormal interactions among brain networks and noted increased information segregation within the brains of schizophrenia patients compared to the healthy control group.

This finding offers valuable insights into schizophrenia. However, it is crucial to recognize that disruptions in integration and segregation are observed across a variety of psychiatric disorders [13]. This broader occurrence reduces the specificity of these biomarkers for schizophrenia. To enhance diagnostic accuracy, future research should examine the extent of overlap in network disruptions across these conditions, with the potential to identify more distinctive patterns or incorporate additional biomarkers. Moreover, longitudinal studies assessing these features at various stages of illness could help determine whether these disruptions are early indicators of schizophrenia or common to other psychiatric disorders.

Additionally, our current study focuses solely on information segregation and integration, without considering their interaction with advancing age. It would be intriguing for us to simultaneously explore both information integration and segregation and how they change with age. This approach could enhance our understanding of the complex relationship between psychosis and the aging process.

Moreover, augmenting the sample size stands out as a pivotal avenue to more comprehensively investigate the psychosis. Additionally, regard to age, it is imperative to factor in the onset of schizophrenia at various ages, as this variable may exert a significant impact on our analytical framework. Lastly, an alternative approach could involve scrutinizing the relationship between functional brain networks within a narrow age window, shedding light on their dynamic changes over a shorter duration. This could not only contribute to a nuanced understanding of age-related dynamics but also offer insights into the potential for timely schizophrenia diagnosis.

Furthermore, the assessment of FCs among various subjects depended on Pearson correlation, overlooking the intricate nonlinear and dynamic characteristics inherent in brain activity [10]–[12]. The omission of considering the human brain as a complex system, particularly in the context of psychotic brain development, underscores the need for additional exploration into dynamic functional connectivity. This approach aims to unravel the nonlinear dynamics associated with the early stages of psychotic brain development. Secondly, the pairwise functional brain connectivity derived from Pearson correlation predominantly considers bivariate connections and overlooks the comprehensive organization of information flow within the entire brain, encompassing interactions beyond pairwise brain regions [34]. A more comprehensive approach, capturing high-order functional connectivity, should be explored to provide a richer understanding of functional connectivity compared to the conventional low-order functional connectivity metrics.

Lastly, regarding the estimation of connections between brain networks and behavior in schizophrenia, it is essential to examine their relationships thoroughly. Changes in interactions among specific brain networks can profoundly influence various behaviors observed in individuals with schizophrenia. Understanding these relationships can offer critical insights into how alterations in brain network dynamics contribute to the manifestation of symptoms and behaviors characteristic of schizophrenia.

## Data and code availability

The preprocessed COBRE dataset can be downloaded from https://figshare.com/articles/dataset/COBRE_reprocessed_with_NIAK_0_12_4/1160600/12.

The corresponding author also can supply all datasets and codes upon a reasonable request.

## Acknowledgements

This work was supported by STI 2030-Major Projects 2022ZD0208900, National Natural Science Foundation of China (No. 62333003, 62036003, 82121003, 62276051, 12271429, 12090021, and 12226007); Medical-Engineering Cooperation Funds from University of Electronic Science and Technology of China

(ZYGX2021YGLH201); China Postdoctoral Science Foundation (Certificate Number: 2024M750360);

## CRediT authorship contribution statement

**Qiang Li**: Writing – review & editing, Writing – original draft, Validation, Methodology, Investigation, Formal analysis, Conceptualization. **Wei Huang**: Writing – review & editing, Methodology, Conceptualization. **Chen Qiao**: Writing – review & editing, Methodology, Conceptualization. **Huafu Chen**: Writing – review & editing, Supervision, Resources, Methodology, Funding acquisition, Conceptualization.

## References

[1] V. Calhoun, J. Sui, K. Kiehl, J. Turner, E. Allen, and G. Pearlson, “Exploring the psychosis functional connectome: Aberrant intrinsic networks in schizophrenia and bipolar disorder,” Frontiers in psychiatry, vol. 2, p. 75, Nov. 2023.

[2] A. C. Yang, C.-J. Hong, Y.-J. Liou, et al., “Decreased resting-state brain activity complexity in schizophrenia characterized by both increased regularity and randomness: Resting brain complexity and schizophrenia,” Human brain mapping, vol. 36, Feb. 2023.

[3] T. Hummer, M. Yung, J. Goñi, et al., “Functional network connectivity in early-stage schizophrenia,” Schizophrenia Research, vol. 218, Feb. 2023.

[4] C. Correll, A. Martin, C. Patel, et al., “Systematic literature review of schizophrenia clinical practice guidelines on acute and maintenance management with antipsychotics,” NPJ schizophrenia, vol. 8, p. 5, Feb. 2023.

[5] J. Duan, M. Xia, F. Womer, et al., “Dynamic changes of functional segregation and integration in vulnerability and resilience to schizophrenia,” Human Brain Mapping, vol. 40, Jan. 2023.

[6] R. Passiatore, L. Antonucci, T. Deramus, et al., “Age-related network connectivity pattern changes are associated with risk for psychosis,” European Psychiatry, vol. 65, S316–S316, Sep. 2023.

[7] M. Heuvel and H. Pol, “Exploring the brain network: A review on resting-state fmri functional connectivity,” European neuropsychopharmacology : the journal of the European College of Neuropsychopharmacology, vol. 20, pp. 519–34, Aug. 2023.

[8] O. Sporns, “Network attributes for segregation and integration in the human brain,” Current opinion in neurobiology, vol. 23, Jan. 2023.

[9] Q. Li, “Functional connectivity inference from fmri data using multivariate information measures,” Neural Networks, vol. 146, pp. 85–97, 2022.

[10] Q. Li, G. V. Steeg, S. Yu, and J. Malo, “Functional connectome of the human brain with total correlation,” Entropy, vol. 24, no. 12, 2022.

[11] Q. Li, V. D. Calhoun, T. D. Pham, and A. Iraji, “Exploring nonlinear dynamics in brain functionality through phase portraits and fuzzy recurrence plots,” Chaos: An Interdisciplinary Journal of Nonlinear Science, vol. 34, Oct. 2023.

[12] Q. Li, G. Ver Steeg, and J. Malo, “Functional connectivity via total correlation: Analytical results in visual areas,” Neurocomputing, p. 127–143, Dec. 2023.

[13] L. D. Lord, A. Stevner, G. Deco, and M. Kringelbach, “Understanding principles of integration and segregation using whole-brain computational connectomics: Implications for neuropsychiatric disorders,” Philosophical Transactions of The Royal Society A Mathematical Physical and Engineering Sciences, vol. 375, p. 20–160 283, May ||2017.

[14] J. Shine, “Neuromodulatory influences on integration and segregation in the brain,” Trends in Cognitive Sciences, vol. 23, May 2019.

[15] P. Fransson and M. Strindberg, “Brain network integration, segregation and quasi-periodic activation and deactivation during tasks and rest,” NeuroImage, vol. 268, p. 119–890, Jan. 2023.

[16] Q. Yu, J. Sui, K. Kiehl, G. Pearlson, and V. Calhoun, “State-related functional integration and functional segregation brain networks in schizophrenia,” Schizophrenia research, vol. 150, Oct. 2023.

[17] Q. Li, V. Calhoun, A. R. Ballem, S. Yu, J. Malo, and A. Iraji, “Aberrant high-order dependencies in schizophrenia resting-state functional MRI networks,” in NeurIPS 2023 workshop: Information-Theoretic Principles in Cognitive Systems, 2023. [Online]. Available: https://openreview.net/forum?id=ZgMRaX02ck.

[18] C. Aine, H. J. Bockholt, J. Bustillo, et al., “Multimodal neuroimaging in schizophrenia: Description and dissemination,” Neuroinformatics, vol. 15, Oct. 2023.

[19] J. Power, A. Cohen, S. Nelson, et al., “Functional network organization of the human brain,” Neuron, vol. 72, pp. 665–78, Nov. 2023.

[20] J. D. A. Relión, D. Kessler, E. Levina, and S. F. Taylor, “Network classification with applications to brain connectomics,” The Annals of Applied Statistics, vol. 13, no. 3, pp. 1648–1677, 2019.

[21] H. Zhang, X. Chen, Y. Zhang, and D. Shen, “Test-retest reliability of “high-order” functional connectivity in young healthy adults,” Frontiers in Neuroscience, vol. 11, 2017.

[22] Y. Zhang, H. Zhang, X. Chen, S.-W. Lee, and D. Shen, “Hybrid high-order functional connectivity networks using resting-state functional mri for mild cognitive impairment diagnosis,” Scientific reports, vol. 7, no. 1, p. 6530, 2017.

[23] M. Carandini and D. J. Heeger, “Normalization as a canonical neural computation,” Nature Reviews Neuroscience, vol. 13, no. 1, pp. 51–62, 2012.

[24] J. Zhang, Y. Barhomi, and T. Serre, “A new biologically inspired color image descriptor,” in European Conference on Computer Vision, 2012. [Online]. Available: https://api.semanticscholar.org/CorpusID:703293.

[25] C. M. Bishop, Pattern Recognition and Machine Learning (Information Science and Statistics). Springer, 2007.

[26] A. Krizhevsky, I. Sutskever, and G. Hinton, “Imagenet classification with deep convolutional neural networks,” Neural Information Processing Systems, vol. 25, Jan. 2023.

[27] C. Seguin, O. Sporns, and A. Zalesky, “Brain network communication: Concepts, models and applications,” Nature Reviews Neuroscience, vol. 24, Jul. 2023.

[28] M. Chan, L. Han, C. Carreno, et al., “Long-term prognosis and educational determinants of brain network decline in older adult individuals,” Nature Aging, vol. 1, Nov. 2023.

[29] D. Chen, Y. Fu, and M. S. Shang, “A fast and efficient heuristic algorithm for detecting community structures in complex networks,” Physica A: Statistical Mechanics and its Applications, vol. 388, pp. 2741–2749, Jul. 2023.

[30] R. Wang, M. Liu, X. Cheng, Y. Wu, A. Hildebrandt, and C. Zhou, “Segregation, integration, and balance of largescale resting brain networks configure different cognitive abilities,” Proceedings of the National Academy of Sciences, vol. 118, e2022288118, Jun. 2023.

[31] D. Bassett, N. Wymbs, M. Porter, P. Mucha, J. Carlson, and S. Grafton, “Dynamic reconfiguration of human brain networks during learning,” Proceedings of the National Academy of Sciences of the United States of America, vol. 108, pp. 7641–6, May 2011.

[32] N. Dosenbach, D. Fair, F. Miezin, et al., “Distinct brain networks for adaptive and stable task control in humans,” Proceedings of the National Academy of Sciences of the United States of America, vol. 104, pp. 11–073–8, Jul. 2023.

[33] X. Wang, Z. Chang, and R. Wang, “Opposite effects of positive and negative symptoms on resting-state brain networks in schizophrenia,” Communications Biology, vol. 6, 2023.

[34] F. Battiston, G. Cencetti, I. Iacopini, et al., “Networks beyond pairwise interactions: Structure and dynamics,” Physics Reports, vol. 874, Jun. 2023.

